# Pan-cancer discovery of driver mutations in long noncoding RNAs reveals widespread functional rewiring of RNA regulatory elements

**DOI:** 10.64898/2026.06.10.730876

**Authors:** Sunandini Ramnarayanan, Roofiya Koya, Marta Soszynska-Jozwiak, Marta Szabat, Jack Roban, Sarang Bhutada, Bhavya Dhaka, Tilman Schaefer, Ai Ming Tham, Hugo A. Guillen Ramirez, Francesca Alessandroni, Michela Coan, Artem Baranovskii, Lambert Moyon, Annalisa Marsico, Colm Ryan, Richard S Houlston, Ryszard Kierzek, Elzbieta Kierzek, Rory Johnson

## Abstract

Most somatic mutations in cancer occur outside protein-coding genes, yet the functional impact of these mutations remains largely unknown. Long noncoding RNAs (lncRNAs) represent a major class of cancer-promoting genes whose molecular mechanisms are poorly understood. While individual driver mutations in lncRNAs have been identified, detecting such ‘driver lncRNAs’ at scale requires large tumour genome cohorts. We analyse 12,631 cancer genomes from the 100,000 Genomes Project (100kGP) and identify 121 lncRNAs under positive selection across 19 cancer types. These driver lncRNAs are independently supported by functional genomic screens, germline predisposing variants, mutual exclusivity with protein-coding drivers, and independent oncogenic lncRNA catalogues. Overall, approximately two-thirds of analysed tumours harbour at least one lncRNA driver mutation. Leveraging the depth of this dataset, we demonstrate that somatic mutations preferentially target and remodel RNA-binding protein (RBP) interaction sites to potentiate oncogenic lncRNAs, including *MALAT1*, *SNHG14* and *NEAT1*. From these data, we derive a model in which somatic mutations liberate oncogenic lncRNAs from repressive RNA:protein interactions. This work expands the number and nature of cancer driver genes, identifies targets for RNA-directed therapies, and demonstrates that with large tumour mutation catalogues we can dissect the molecular mechanisms of noncoding genes.

## Introduction

Cancer initiation and progression are driven by positively selected somatic mutations that confer a cellular growth advantage^1^. While protein-coding driver mutations have been extensively catalogued^2,3^, over 90% of somatic mutations occur outside protein-coding regions^1^. The functional impact of these non-coding mutations remain largely unclear, however multiple lines of evidence support the existence of numerous hitherto undiscovered fitness-altering driver mutations^4–17^. Previous pan-cancer efforts, including the Pan-Cancer Analysis of Whole Genomes (PCAWG) consortium, confirmed non-coding drivers exist but were limited by cohorts of fewer than 3,000 genomes, which reduces statistical power to detect rare or weak-effect drivers^12^.

Fitness-altering somatic mutations in non-coding RNAs have nevertheless been reported. In addition to well-established regulatory *TERT* promoter mutations^18,19^, recurrent somatic mutations in small non-coding RNAs, such as the U1 spliceosomal RNA^20^, have been shown to disrupt RNA–protein interactions and splicing. These findings demonstrate that somatic mutations can target structural and functional elements of non-coding RNAs to promote tumourigenesis.

Long noncoding RNAs (lncRNAs) constitute the most numerous and functionally diverse class of non-coding genome transcripts^21,22^. Hundreds of lncRNAs are dysregulated in cancer and can promote tumourigenesis *in vitro* and *in vivo*^23,24^. LncRNAs are thought to function through modular assemblies of discrete elements, including conserved secondary structures and docking sites for RNA-binding proteins (RBPs)^25^. We have recently provided the first systematic evidence that lncRNAs can act as cancer drivers^21^. In particular, based on a limited analysis of 3000 tumours and experimental validation, we have demonstrated that mutations in *NEAT1* can remodel its ribonucleoprotein complex to promote paraspeckle formation, resulting in accelerated cell and tumour growth^22^.

To systematically define the landscape and mechanisms of lncRNA drivers in cancer, we here analyse the largest tumour genome collection to date, assembled by the 100,000 Genomes Project (100kGP) and comprising whole-genome sequences (WGS) from 12,631 tumours. Using a new population-scale driver discovery pipeline, we identify 121 lncRNAs under positive selection. We validate their disease relevance by reference to independent functional and clinical datasets. Crucially, the scale of the 100kGP cohort enables us to link driver mutations directly to mechanistic features, revealing a model in which somatic mutations rewire functional elements of oncogenic lncRNA to drive cancer.

## Results

### Tumour genomes from the 100,000 Genomes Project (100kGP) enable large-scale discovery of cancer driver lncRNAs

To generate a comprehensive map of mutated cancer driver lncRNAs (‘driver lncRNAs’), we analysed WGS data from the 100kGP cohort (Fig. 1a). We began with an initial dataset of 16,341 tumour genomes and 573 million small somatic variants. To establish a high-confidence dataset for downstream analysis, we applied a rigorous quality control pipeline that included sample deduplication, the removal of microsatellite instability cases, and the exclusion of tumours lacking sufficient corroboration between genomic and secondary clinical data (see Methods)^17,26^. We further required that a cancer type needed to have at least 20 samples and restricted variants to single-nucleotide variants and 1 bp indels lying outside of regions prone to sequencing artefacts (collectively ‘SNVs’)^27,28^. These filters yielded a final high-quality dataset of 12,631 tumour genomes and 238,807,909 SNVs across 37 cancer types (Fig. 1b). Numbers of Breast (BRCA), Colon (COAD), Kidney (KIRC), and Sarcoma (SARC) tumours were particularly large, each exceeding 1,000 samples (Fig. 1c). Power calculations showed cohorts of 13 cancer types being sufficiently large to identify lncRNAs mutated at ≥5% above background with 90% confidence (Fig. 1d), compared with only six in PCAWG^4^.

**Figure 1:**
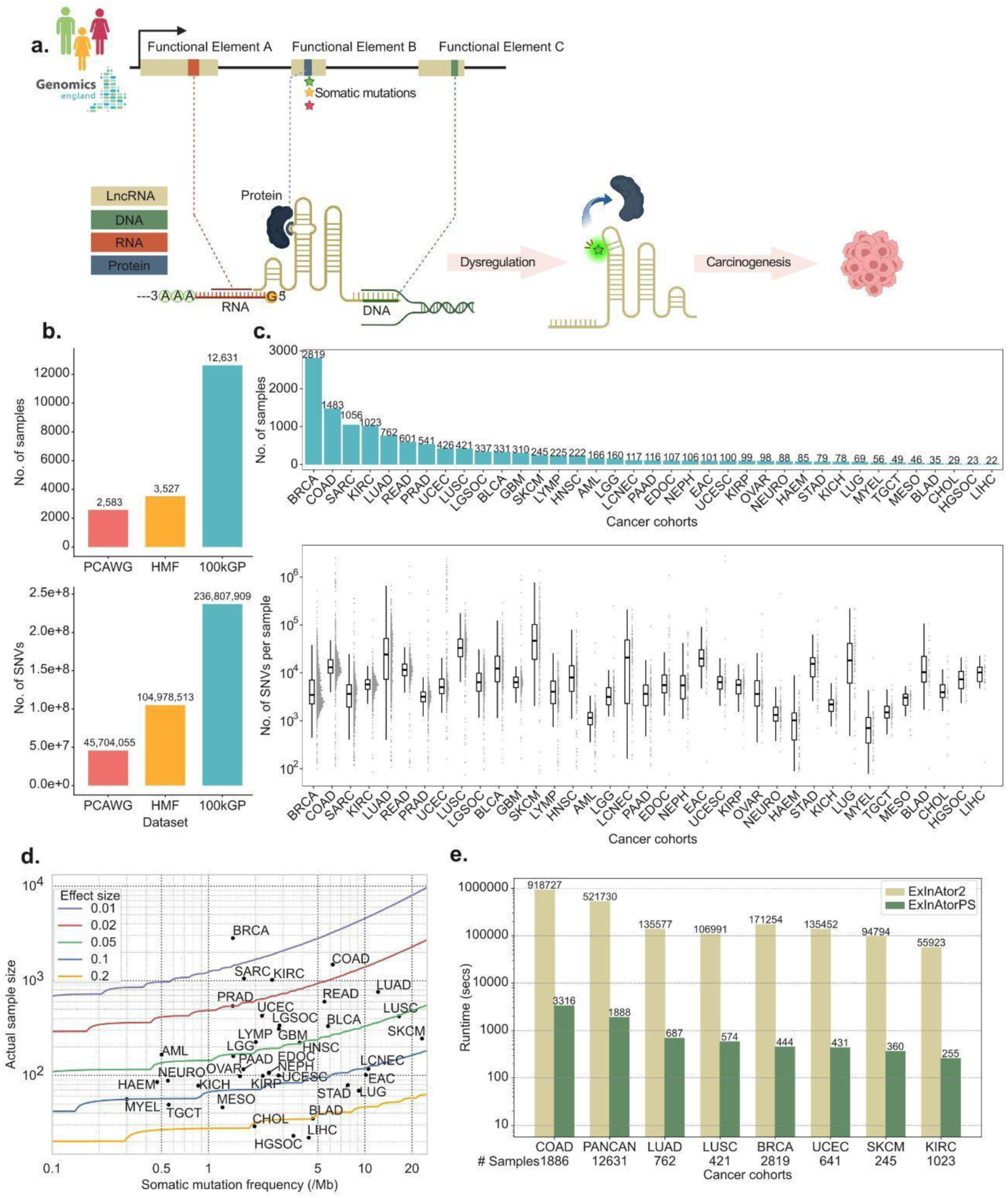
Large-scale discovery of cancer driver lncRNAs in the 100,000 Genomes Project (100kGP) **a.** Model showing how somatic mutations in lncRNAs can increase cellular fitness and drive carcinogenesis through alterations to functional elements. **b.** Number of tumour samples (top) and single-nucleotide variants (SNVs, bottom) across three datasets: 100kGP, HMF (Hartwig Medical Foundation) and PCAWG. **c.** Sample size (upper) and total number of SNVs (lower) for each cancer type in the 100kGP tumour collection. **d.** Power analysis showing the minimum number of samples required to detect driver lncRNAs at 90% power for different effect sizes (foreground mutation rate relative to background). **e.** Runtime comparison for ExInAtor2 and ExInAtorPS driver discovery pipelines.

Our goal was to detect driver lncRNAs using the published ExInAtor2 pipeline, which searches for established signatures of positive selection: mutational burden and functional impact^27,28^. ExInAtor2 works on the principle that lncRNA driver mutations act by modifying the mature RNA sequence, and hence will fall in exonic regions. Although the existing ExInAtor2 pipeline had successfully detected drivers in the smaller PCAWG dataset (∼2,500 tumour genomes)^21^, it proved computationally intensive for the 100kGP tumour collection, with run times of 1-10 days per cancer type (Fig. 1e). We therefore developed ExInAtorPS (population scale), featuring accelerated empirical significance estimation to enable analysis of population-scale cohorts without loss of accuracy (see Methods). Compared to ExInAtor2, ExInAtorPS analysed the full 100kGP pan-cancer dataset ∼276-fold faster (in ∼31 minutes) and with an average 265-fold improvement across cancer types (Fig. 1e).

### High accuracy and low false discovery rate of driver lncRNA discovery

We first validated ExInAtorPS predictions in the 100kGP cohort. Since ExInAtorPS is agnostic to gene biotype, we benchmarked its performance on protein-coding genes (PCGs) using the COSMIC Cancer Gene Census (CGC) as ground truth^29^. When run on all 37 individual cancer types, as well as their union (Pan-Cancer, ‘PANCAN’), ExInAtorPS identified 421 PCG drivers, achieving a median precision of 67.4% (FDR≤10%; Fig. S1.a). Precision increased to 80% when compared to PCG drivers from IntOGen, TCGA and 100kGP^2,30,31^. All except Myeloma (MYEL) showed significant enrichment of true positives (overall 17.8-fold enrichment, P=9e-109, one-sided Fisher’s exact test) with only 3.3% (14 genes) overlapping known false positives (Fig. S1.b, S1.c, S1.d)^32,33^.

Having established the accuracy of ExInAtorPS, we applied it to lncRNAs. We supplemented intergenic lncRNAs from GENCODE^34^ with 648 experimentally-validated cancer lncRNAs from the Cancer LncRNA Census 3 (CLC3) dataset, yielding a test set of 7,794 filtered lncRNA genes^35^ (see Methods).

In the 100kGP dataset, ExInAtorPS identified 172 driver lncRNA-tumour pairs, corresponding to 122 drivers across 18 cancer types and PANCAN (FDR ≤10%, Fig. 2a). The highest numbers of driver lncRNAs were observed in Breast (BRCA, 27), Bladder (BLCA, 25), Melanoma (SKCM, 21), Lung (LUAD, 11) and the pan-cancer analysis (PANCAN, 58). No drivers were identified in the remaining 19 cancer types, 10 of which had small numbers of cases (<100). Post-candidate filtering led to 166 drivers, corresponding to 121 unique lncRNAs across 18 cancer types and PANCAN (see Methods).

**Figure 2:**
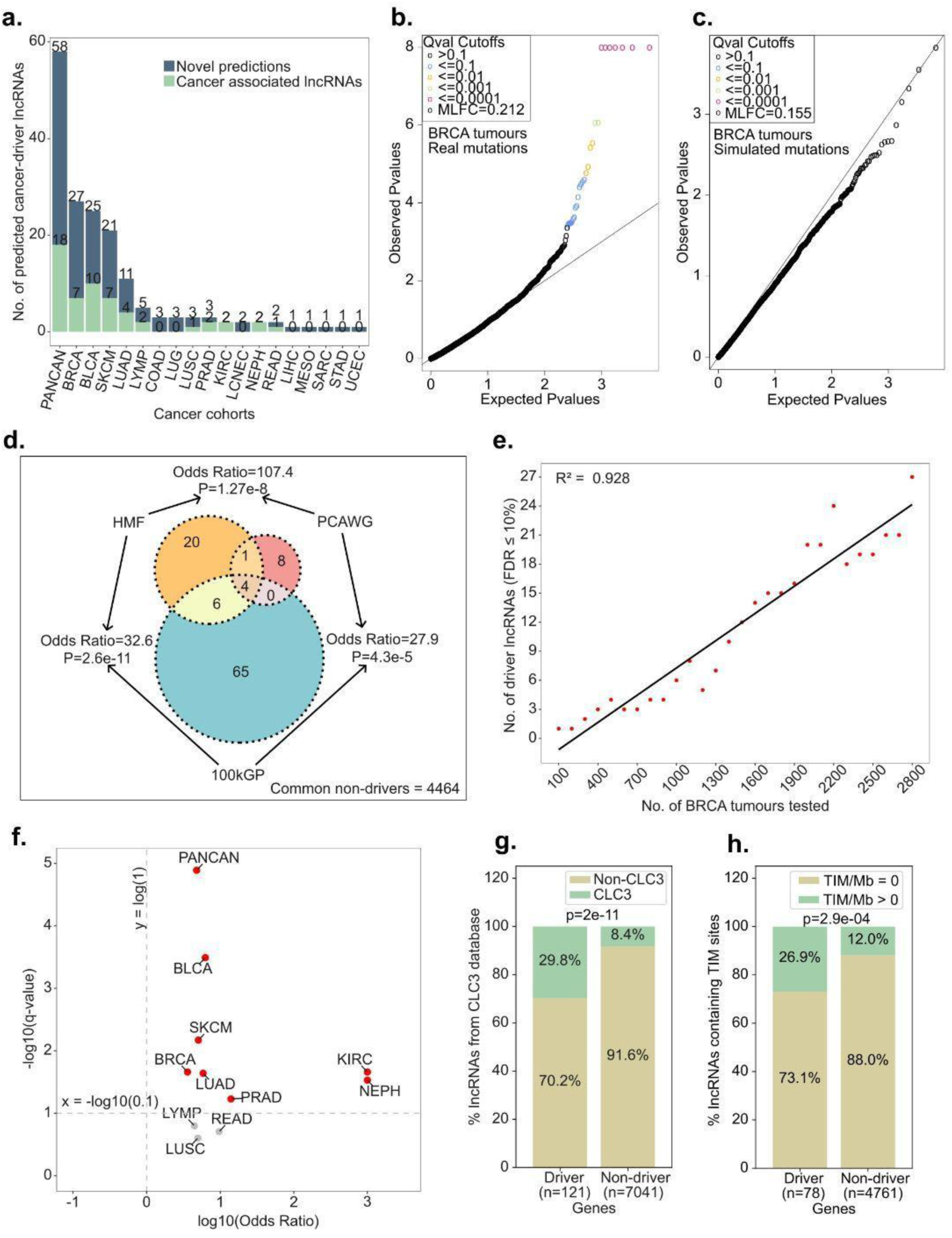
ExInAtorPS identifies driver lncRNAs with high specificity and independent support. **a.** Number of predicted driver lncRNAs across cancers. Green and blue bars indicate lncRNAs that do or do not overlap, respectively, known cancer-associated lncRNAs from the Cancer LncRNA Census 3 (CLC3) database. The upper number denotes the total number of driver predictions, while the lower number denotes the number of driver predictions that also belong to the CLC3 set. **b.** Quantile-Quantile (QQ) plot of observed vs expected p-values for real mutations in BRCA cases. Mean log fold change (MLFC) measures deviation from the expected neutral distribution, where values near zero indicate a well-calibrated model. **c**. As for (b), but for simulated mutations in BRCA cases (no drivers identified at FDR≤10%)**. d.** Overlap of driver lncRNAs identified in 100kGP, HMF and PCAWG WGS datasets. **e.** Downsampling analysis showing the number of driver lncRNAs detected as a function of BRCA sample size. **f.** Overrepresentation of CLC3 genes among predicted driver lncRNAs across cancer cohorts. **g.** Enrichment of CLC3 genes in driver vs non-driver lncRNAs by cancer type. **h.** Enrichment of transposon insertional mutagenesis (TIM) insertion sites (TIM/Mb) in driver versus non-driver lncRNAs.

Quantile-quantile (QQ) plots confirmed that most lncRNAs followed the expected null distribution (Fig. 2b). ExInAtorPS conservatively corrects for mutational signatures using local trinucleotide rate correction, as employed previously for ExInAtor2^21,22^. However, to formally test the false discovery rate, we conducted permutation testing (100 randomised SNV sets) on the BRCA cohort, which yielded an average of only 0.15 false discoveries (Fig. 2c, S1.e, S1.f), equivalent to an empirical FDR of 0.6% vs ExInAtorPS nominal 10%, thereby demonstrating high specificity in driver identification (see Methods).

Reproducibility was assessed by comparing 100kGP driver lncRNAs to those identified previously in the PCAWG and Hartwig Medical Foundation (HMF) cohorts^21^ (Fig. 2d). The overlap between each pair of projects is statistically significant but relatively modest, consistent with the notion that sensitivity to detect driver lncRNAs is constrained by genome collection size, and compounded with the fact that these cohorts focus on different cancer types and patient age ranges. Downsampling analysis of BRCA tumours showed that the number of detected driver lncRNAs increased linearly with the cohort size (R^2^=0.93, Fig. 2e) and did not plateau, indicating that many true driver lncRNAs probably remained undiscovered.

When considering driver lncRNAs from each cancer type separately, they were significantly enriched for literature-curated cancer genes from the ground truth CLC3 set in all but three cases (Fig. 2f), with 29.8% (36/121) of all driver lncRNAs overlapping these genes^35^ (4.6-fold enrichment, P=2e-11, one-sided Fisher’s exact test, Fig. 2g). Driver lncRNAs were 2.7-fold enriched for driver genes discovered by published experimental transposon insertional mutagenesis (TIM)^35–37^(P=2.9e–4, one-sided Fisher’s exact test) (Fig. 2h). In sum, multiple lines of orthogonal evidence support the disease relevance of predicted driver lncRNAs.

### Pan-cancer landscape of cancer driver lncRNAs

The 121 candidate driver lncRNAs were frequently mutated across the individual tumours of the 100kGP (Fig. 3a). Mutations in the known oncogenic lncRNA *SNHG14* are the most frequently observed, occurring in 4,390 tumours (35%)^38^. Of the 121 driver lncRNAs, 12 were identified in ≥2 cancer types, and 7 in ≥3 cancer types (Fig. 3b). *NEAT1,* the first experimentally validated driver lncRNA, is identified in the greatest number of cancer types (4)^21^. The well-established cancer-associated lncRNAs *RMRP* and *MALAT1*, both of which had been reported as mutated drivers by PCAWG^11,17,39,40^, were the next most recurrent in the 100kGP dataset. Many driver lncRNAs showed a mutational burden several folds above background (Fig. 3b). For example, the previously reported oncogene *LINC00324* exhibits a 2.8-fold and 1.8-fold excess of exonic mutations in Breast and PANCAN cohorts, respectively^41,42^.

**Figure 3:**
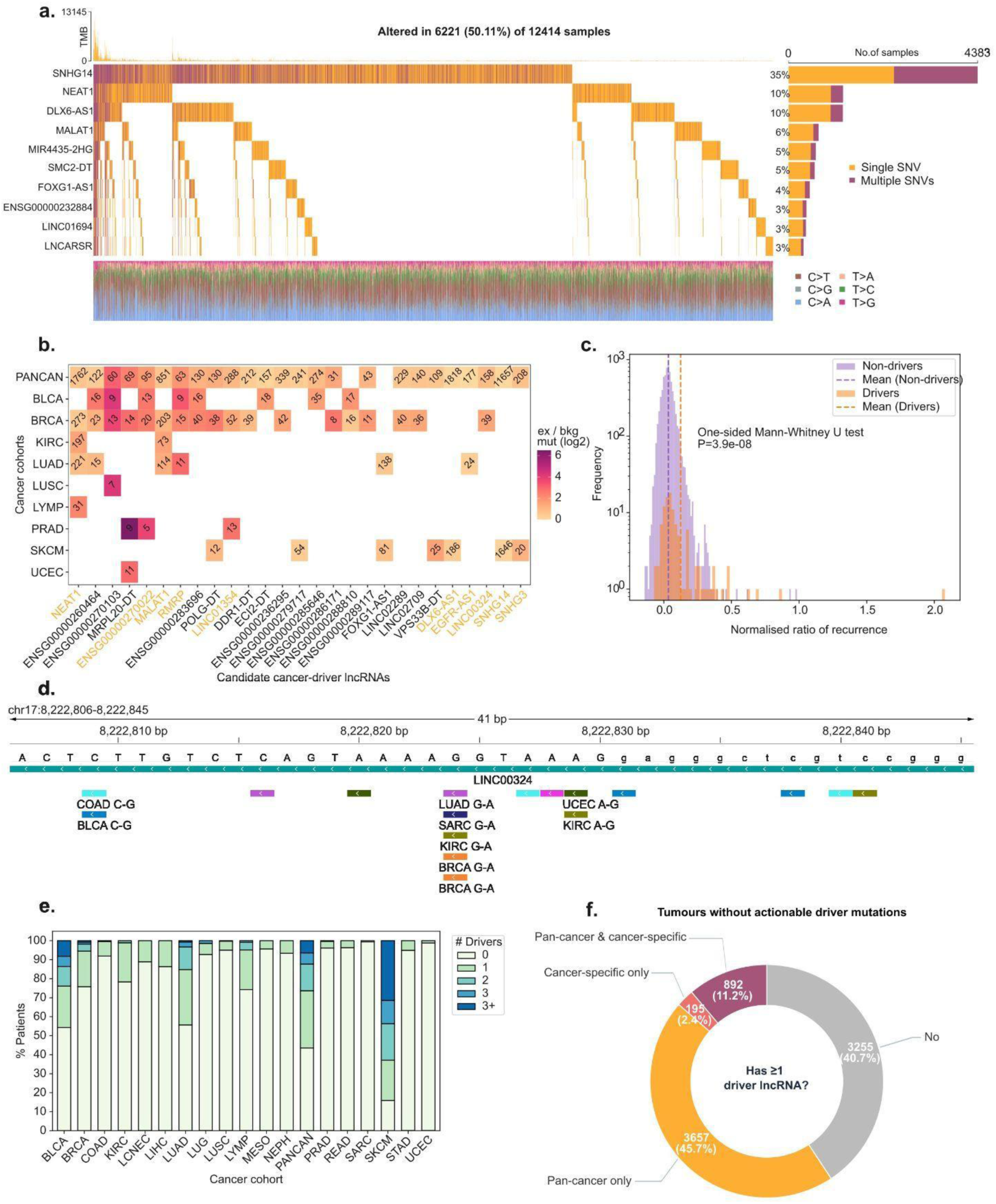
Landscape of driver lncRNAs across 12,631 cancers. a. Mutations in driver lncRNAs across tumour samples. Columns represent tumours, where orange and purple indicate single and multiple SNVs per patient, respectively. b. Mutation burden in driver lncRNAs across multiple cancer types (only driver lncRNAs identified in ≥2 cancers are shown). Cell colour indicates the fold enrichment of exonic over background mutation rate; numbers show count of exonic mutations. c. Normalised recurrence ratio of mutations in driver and non-driver lncRNAs, adjusted for length and trinucleotide context. D. A region of *LINC00324* driver lncRNA, containing SNVs (coloured rectangles) that recur across patients and cancer types. e. Proportion of tumours with mutations in driver lncRNAs. Bars indicate the fraction of tumours harbouring 0, 1, 2, or 3+ cohort-specific mutated driver lncRNAs. f. Mutated driver lncRNAs in tumours without approved targeted therapy.

As evidenced by promoter mutations in *FOXA1* and *TERT*, recurrent mutations at the same nucleotide position provide strong evidence for positive selection^11,18,19,43^. SNVs in driver lncRNAs were significantly more recurrent than expected, even after adjusting for mutational burden, length and signature (Fig. 3c). *LINC00324* additionally harbours recurrent alterations across cancer types, consistent with positive selection (Fig. 3d).

Driver lncRNAs represent promising therapeutic targets for precision oncology^39,44^. Across the entire cohort of 12,631 tumour genomes, 7,239 (54%) harboured at least one driver mutation in a cohort-specific lncRNA (ie a lncRNA predicted to be a driver in that same cancer type); this increased to 8,206 (65%) when considering any identified driver lncRNAs regardless of cancer type. Equivalent rates in bladder urothelial carcinoma (BLCA) and in lung adenocarcinoma (LUAD) exceeded 40%, whereas it was over 80% in skin cutaneous melanoma (SKCM) (Fig. 3e). The latter may be partly attributed to hypermutation. Among the 7,999 tumours lacking approved targeted therapy, 59% (4,744) carried at least one driver lncRNA mutation, potentially representing a therapeutic target (Fig. 3f)^31^.

### Clinico-biological characteristics of driver lncRNAs

Driver lncRNA genes exhibited properties expected of biologically functional genes^45^: they were expressed higher in the healthy tissue of origin for their cancer type (Fig. 4a) and were expressed in tumours where they were mutated (Fig. 4b), consistent with expression preceding mutation as expected for genuine driver genes.

**Figure 4:**
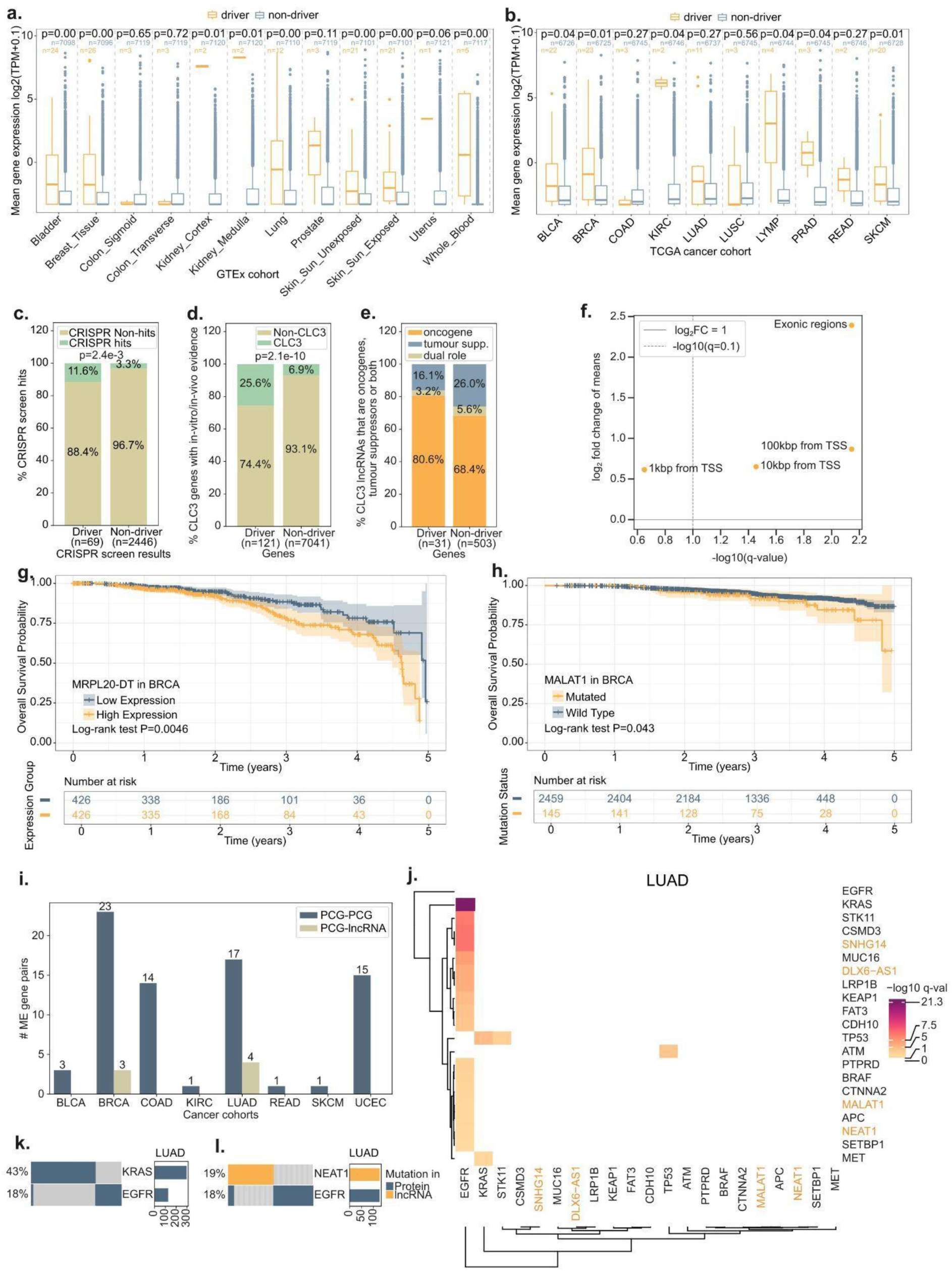
Clinico-biological characteristics of driver lncRNAs. a. Driver lncRNAs and non-driver lncRNA expression levels from RNA-sequencing of matched healthy organs. For each tissue type, only driver lncRNAs from that organ’s corresponding cancer are considered. b. As for (a), but in tumours. c. Intersection of driver lncRNAs with fitness-affecting hits from CRISPR viability screens (P=2.4e-3, one-sided Fisher’s exact test). d. Intersection of driver lncRNAs with lncRNAs that were previously shown to be disease-modifying in the CLC3 resource (P=2.1e-10, one-sided Fisher’s exact test). e. The proportion of oncogenes, tumour-suppressors, and dual-role genes in driver and non-driver lncRNAs, as annotated by the CLC3 resource. f. Enrichment of germline cancer-predisposing SNPs from the EBI-GWAS catalogue near driver lncRNAs relative to non-drivers across different proximity windows. g. Kaplan–Meier survival analysis in BRCA stratified by MRPL20-DT expression. LncRNAs were equally divided into high and low expression groups. h. Kaplan–Meier survival analysis stratified by *MALAT1* mutation status (mutated vs wild-type). i. Numbers of significantly mutually exclusive (ME) gene pairs involving lncRNAs across different cancers. j. Mutually exclusive gene pairs in LUAD. Orange: driver lncRNAs. k. LUAD tumours carrying *KRAS* and *EGFR* mutations in LUAD. l. LUAD tumours carrying *NEAT1* and *EGFR* mutations in LUAD.

We explored whether driver lncRNAs had disease-modifying potential. We interrogated published CRISPR viability screen datasets^23,24^ and found that driver lncRNAs were significantly enriched among fitness-impacting hits across multiple cancer cell lines (Odds ratio = 3.8, P=2.4e-3, one-sided Fisher’s exact test) (Fig. 4c). We also found that driver lncRNAs were enriched for the subset of CLC3 genes previously shown to be disease-modifying by *in vitro* or *in vivo assays* (Odds ratio = 4.68, P=2.1e-10, one-sided Fisher’s exact test) (Fig. 4d). Although not statistically significant, an enrichment of oncogenes over tumour-suppressors (as defined in CLC3) was observed amongst drivers lncRNAs (Odds Ratio = 1.93, P=0.11, one-sided Fisher’s exact test) (Fig. 4e).

Somatically-mutated driver genes often harbour germline cancer susceptibility variants^46^. We compared the density of known cancer-associated single nucleotide polymorphisms (SNPs) from the EBI-GWAS catalogue in driver and non-driver lncRNAs, within exons or within various ranges of the transcription start site (TSS). Driver lncRNAs were significantly enriched for susceptibility SNPs in exonic regions (Fig. 4f) (P-adj = 0.007) and within 10 kb (P-adj=0.035) and 100 kb of the TSS (P-adj =0.007, BH-adjusted, one-sided Fisher’s exact test) (Fig. 4f).

We next evaluated the prognostic effect of driver lncRNA expression levels within each cancer type. Since gene expression data is not available for 100kGP, we used cancer-matched TCGA data. After correcting for multiple testing, the expression of three driver lncRNAs was significantly associated with patient survival (FDR≤5%). For example, higher levels of BRCA-driver MRPL20-DT was associated with poorer patient survival, remaining significant after adjusting for age and tumour stage (Fig. 4g, Fig. S2a). We also searched for prognostic effects of driver lncRNA mutations, but could identify no cases that remained significant after multiple testing correction within each cancer type. Patients with *MALAT1* mutations showed reduced survival, although this association did not remain significant after adjustment (Fig. 4h, Fig. S2b).

Driver gene mutations often display mutually exclusive relationships, which can result from functional linkages between those genes^47,48^. In the 100kGP cohort we correctly identified known pairs of mutually exclusive protein-coding mutations, linking *EGFR* and *KRAS* in Lung (LUAD) (Fig. 4i, 4j, 4k), and *BRAF* and *NRAS* in Melanoma (SKCM, Supplementary File 4)^49,50^. Next, we extended this analysis to include driver lncRNAs and we identified seven mutually exclusive pairs involving five lncRNAs and four protein-coding genes at FDR≤1% (Fig. 4i). These comprise *NEAT1* / *EGFR*, *DLX6-AS1 / EGFR, SNHG14 / EGFR, MALAT1 / EGFR in LUAD (Fig. 4j, 4l) and MIR4453-2HG / PIK3CA, SNHG14 / CBFB* and *SNHG14 / NCOR1* in BRCA (Fig. S2d). The mutual exclusivity between mutations in *NEAT1* and *EGFR* is consistent with the fact that NEAT1 is regulated by EGFR signalling^51^ (Fig. 4l). In contrast to mutual exclusivity, mutational co-occurrence can reflect synergy between pairs of mutations. We also performed co-occurrence analysis, and discovered five lncRNA-protein-coding gene interactions for COAD at FDR≤1% (Fig. S2c, Supplementary File 4). These mutational co-occurrence relationships provide evidence linking driver lncRNA to established oncogenic pathways.

### Driver lncRNA mutations target protein binding sites and secondary structures

We had previously shown that fitness-enhancing mutations in *NEAT1* altered RNA-binding proteins (RBPs) recruitment^21,25^. This prompted us to hypothesise that driver mutations could act by altering lncRNA functional elements such as RPB binding sites and secondary structures.

To systematically test this, we developed ExInAtorEL, an element-level driver discovery method assessing mutational burden in annotated lncRNA elements relative to trinucleotide-matched background exon sequences (Methods) (Fig. 5a). We curated 250 element classes, including eCLIP-defined RBP binding sites, high-confidence evolutionarily-conserved RNA secondary structures (CRS), and evolutionarily conserved sequence elements (PhastCons, considering 7-, 20- and 100-way phylogenies) (Fig. 5b; see Supplementary File 5)^52–54^.

**Figure 5:**
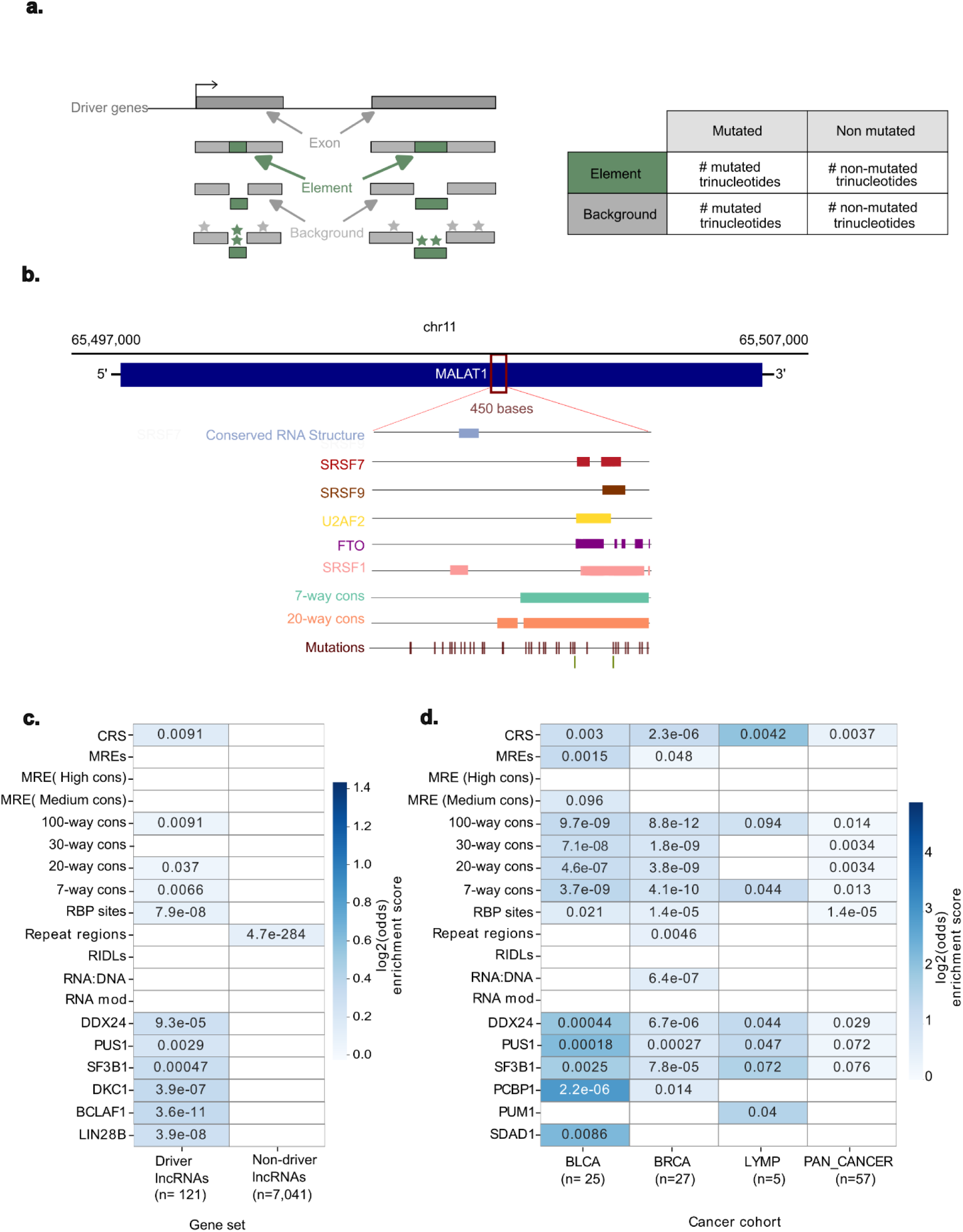
Mutations in driver lncRNAs target diverse RNA functional elements. a. Overview of the ExInAtorEL approach to discover recurrently mutated lncRNA functional elements. Functional elements and background regions are defined within lncRNA exons. Background regions are subsampled to match the trinucleotide composition of each element, enabling unbiased comparison of mutation rates. Mutation enrichment is assessed using one-sided Fisher’s exact test. b. Browser view of a 450-nucleotide region in the *MALAT1* locus showing functional elements: conserved RNA secondary structures (CRS), binding sites for RNA-binding proteins (SRSF7, SRSF9, U2AF2, FTO, and SRSF1), and evolutionarily conserved elements (7-way and 20-way vertebrate PhastCons elements). c. The results for ExInAtorEL analysis of the union of pan-cancer mutations in combination with all 121 driver lncRNAs. Rows contain classes of elements tested, columns contain results for driver lncRNAs or for the remaining tested non-driver lncRNAs. RBP sites were tested both as the union of all sites (‘RBP sites’) and individual RBP types, of which a selection is displayed. The numbers inside cells indicate False Discovery Rate, colour represents degree of mutational enrichment in elements compared to exonic background according to the indicated scale. d. As for (c), except ExInAtorEL was applied to individual cancer cohorts, considering only driver lncRNAs identified from those same cohorts.

When considering the union of pan-cancer mutations and all 121 driver lncRNAs, ExInAtorEL identified significant mutational enrichment in CRS (P = 0.0091), RBP sites (P = 7.9e-08) and PhastCons conserved elements (P = 0.0066, P = 0.037, P = 0.0091 for 7-way, 20-way and 100-way, respectively) (Fig. 5c; one-sided Fisher’s exact test, BH-adjusted, FDR < 10%). Enrichment ranged from 1.04- to 1.5-fold, compared to non-element sequences, and was observed consistently across different cancer types (1.4- to 4-fold; Fig. 5d; Supplementary File 6). Overall, 17,703 SNVs were detected in altogether 191.5 kb of elements within driver lncRNAs (92.4 SNVs / kb), as compared with 398,864 SNVs in 4,715.7 kb of non-driver lncRNAs (84.6 SNVs / kb), corresponding to a global 1.09-fold enrichment (P =8.95e-33, one-sided Fisher’s exact test; Fig. S3a). Annotated elements comprised only a small fraction of exonic sequence (Fig. S3b). Permutation testing on simulated neutral SNVs in BRCA yielded an empirical FDR of 0.2% (0.32 false positives on average versus 158 observed driver elements), supporting the specificity of ExInAtorEL predictions (Fig. S3c) (see Methods).

Several further observations support the validity of driver element predictions. Firstly, mutational enrichment was only observed in driver lncRNAs, but not in non-driver lncRNAs (Fig. 5c). Secondly, the same elements were mutated in different cancers (Fig. 5d) and in driver lncRNAs from the independent PCAWG dataset (Fig. S3d). Conversely, this enrichment signal disappeared when simulated neutral SNVs were analysed (Fig. S3e).

We further tested whether interaction sites of specific RBPs were preferentially targeted by mutations. Of the 150 RBPs in our dataset^52,55^, binding sites for 99 (66%) showed significant mutational enrichment at an FDR < 10%, when considering the union of pan-cancer mutations and all 121 driver lncRNAs (Supplementary File 6). Again, enrichment patterns were consistent across different cancers (Fig. 5d, Supplementary File 6). One example is the RBP SRSF1, which binds to the oncogenic lncRNA MALAT1 and has been linked to tumourigenesis^56–58^. SRSF1 binding sites were significantly mutated across pan-cancer drivers as well as in BLCA and BRCA cohorts highlighting the role of disrupted RBP–lncRNA interactions in cancer (Supplementary File 6).

### Driver mutations alter the activity of lncRNA functional elements

Elevated mutational burden in conserved secondary structures and RBP sites suggests that driver SNVs alter lncRNA function to confer a fitness advantage, for example by changing RNA folding or RBP affinity. Indeed, we observed frequent SNVs in conserved structural regions of driver lncRNAs, such as the oncogenic lncRNA SNHG14^35^, which carried a high density of patient-derived SNVs concentrated within a specific conserved secondary structure region (CRS M0396124) (Fig. 6a). A similar pattern is seen with *EXOC3-AS1,* a novel driver candidate (Fig. S4a).

**Figure 6:**
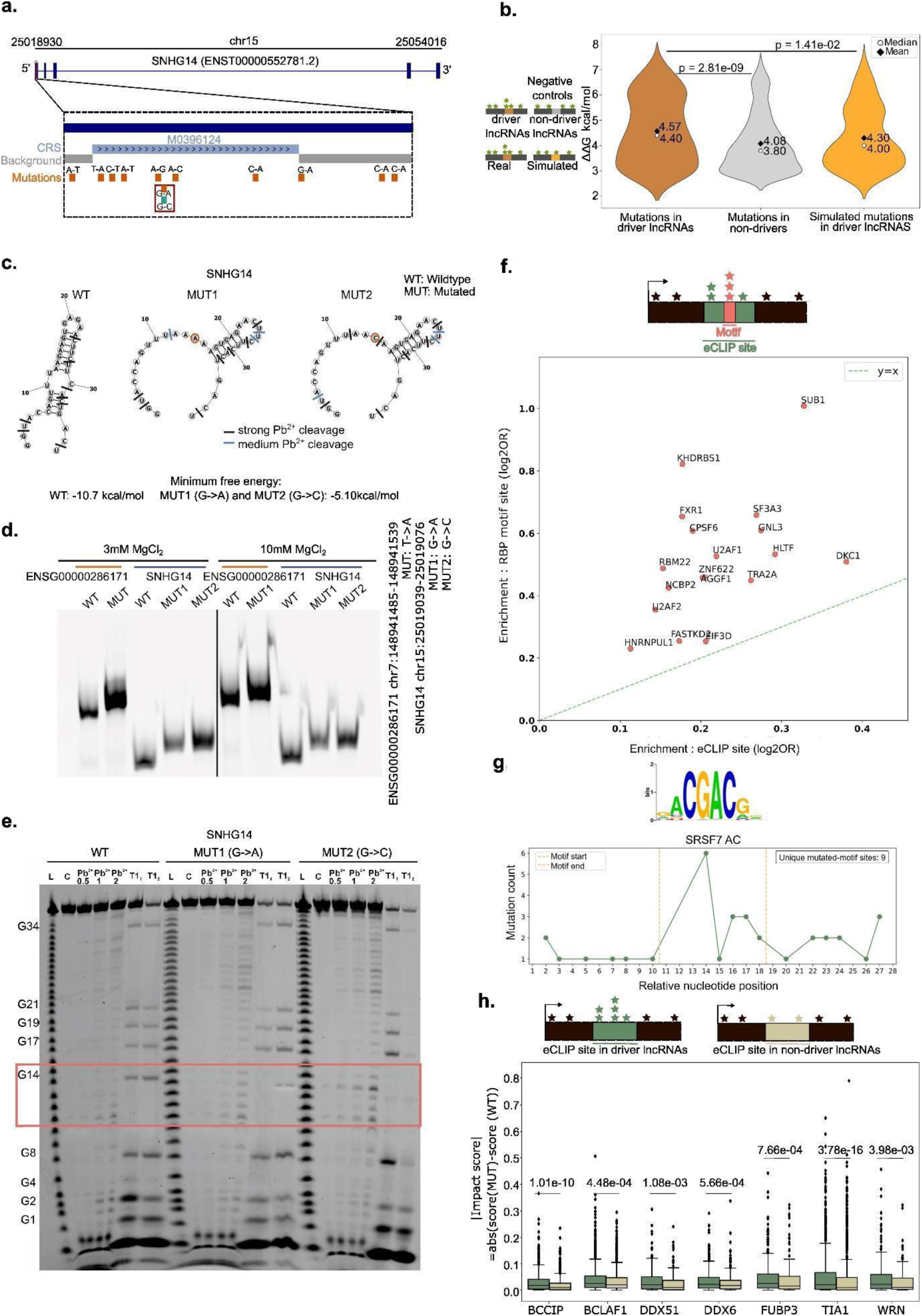
Functional impact of driver mutations on lncRNA elements. a. SNVs in the evolutionarily conserved RNA structure M0396124 in SNHG14. A recurrently mutated position is indicated by the red box. b. Absolute change in predicted RNA folding energy (|ΔΔG|, kcal mol⁻¹) induced by SNVs within CRS elements. Columns contain: Pan-cancer SNVs in the entire set of driver lncRNAs; Pan-cancer SNVs in the non-driver lncRNAs; randomised, trinucleotide-matched pan-cancer SNVs in the driver lncRNAs. Statistical significance was evaluated by one-sided Wilcoxon rank-sum test (alternative = ‘greater’). c. RNAfold-predicted secondary structures of wild-type and mutant conserved RNA structure in SNHG14. The nucleotides accessible to lead ion (Pb²⁺) cleavage are indicated. d. Native gel electrophoresis of wild-type (WT) and mutant (MUT) CRS sequences. The mutations are located in the following genomic locations (assembly GRCh38): ENSG00000286171 - chr7:148941485-148941539(+); SNHG14 - chr15:25019039-25019076(+) e. Lead ion (Pb²⁺) probing of wild-type and mutant CRS in SNHG14. Red box highlights positions 10-14, where mutation-induced gain in Pb^2+^ sensitivity is observed and corresponds to the RNA-fold prediction of loss of base-pairing in (c). f. Mutational enrichment of pan-cancer SNVs at RBP eCLIP-defined binding sites (*x* axis) versus sequence-motif sites within those binding regions (*y* axis), in the entire set of driver lncRNAs. Each point represents the set of binding sites for the indicated RBP. Only RBPs with significant enrichment in both tests are shown (eCLIP sites and motif sites, each FDR < 10%, tested independently). Dashed line, y = x (equal enrichment in both). g. Distribution of pan-cancer SNVs across the SRSF7 motif and flanking sequence, pooled across all driver lncRNAs. The *x* axis indicates the relative nucleotide position in the core motif and ±10 bp of flanking sequence; the *y* axis denotes the number of SNVs at each position. All SNVs falling within SRSF7 motif instances are included. Motif instances within eCLIP peaks were identified from position weight matrices (CISBP-RNA, RBPDB) using MOODS v1.9.4.1 (DNA mode, P < 0.005). h. Predicted impact of pan-cancer SNVs on binding affinity, for SNVs within eCLIP peaks in the entire set of driver lncRNAs versus non-driver lncRNAs. For each RBP (*x* axis), the change in Pysster-predicted RBP-binding probability between the mutated and wild-type alleles- |impact score| (*y* axis) is shown; one-sided Mann–Whitney U test (driver versus non-driver), (FDR < 10%).

Using RNAFold, we calculated the change in minimum free energy (ΔMFE) for CRS elements as a consequence of each driver lncRNA SNV^59^. Most (75%) mutations result in structural destabilisation (Fig. S4b) and were associated with substantially larger changes in folding energy (|ΔMFE|) than SNVs in nondriver lncRNAs (P = 2.81e-09 Mann–Whitney U test; Fig. 6b) or simulated neutral SNVs in driver lncRNAs (P = 1.41e-02 Mann–Whitney U test; Fig. 6b). Mutations with |ΔMFE| > 2.5 kcal/mol were markedly enriched among SNVs of driver lncRNAs (Fig. 6b), consistent with positive selection on alterations of RNA structure.

To add confidence to these *in silico* predictions of RNA structures, we shortlisted and experimentally tested eight candidate SNVs, which confirmed the predictions for five of them (Supplementary File 7). Shortlisting was based on a combination of predicted structural impact, mutational recurrence, and prior published cancer roles for the host lncRNA. In *SNHG14*, two independent SNVs disrupted a hairpin structure, shifting it toward a less stable configuration^60^ (Fig. 6c). Structural rearrangement was also predicted for *LINC01718 and LINC01411* (Fig. S5a,b). Native gel electrophoresis of *in vitro* synthesised RNA confirmed mutation-associated structural changes in *SNHG14, ENSG00000286171, LINC01718,* and *LINC01411* (Fig. 6d and Fig. S5c,d). Lead-ion (Pb2+) probing further demonstrated substantial structural RNA rearrangements in three candidates (Fig. 6e and Fig. S5e,f)^61,62^.

We next assessed the impact of SNVs on RBP binding sites. RBP interactions are mediated by a core recognition motif that underpins molecular recognition, which sits within a broader region that is experimentally defined by eCLIP. We hypothesised that functionally relevant SNVs would be enriched in core motifs, compared to flanking eCLIP regions. Restricting analysis to RBPs with defined recognition motifs (n = 107/150), we observed significant enrichment of mutations within motifs across pan-cancer drivers as well as in BRCA and BLCA (BH-adjusted FDR < 10%; Fig. 6f and Fig. S6a,b), but not in non-driver lncRNAs (Fig. S6e). Notably, mutation density frequently mirrored nucleotide-level importance within motifs (Fig. 6g and Fig. S6c,d).

To gain further insights into the impact of RBP mutations, we turned to a convolutional neural network (CNN) model of RBP binding to quantify the impact of driver SNVs^63^. Compared to mutations in the corresponding eCLIP sites in non-driver lncRNAs, mutations in driver lncRNAs more significantly impacted the predicted RBP:RNA affinity of several cancer-associated RBPs (Mann–Whitney U test, BH-adjusted, FDR < 10%; Fig. 6h). The majority of such changes decreased predicted affinity, although similar to CRS above, there are numerous cases where driver mutations were predicted to increase affinity (Fig. S6g).

Collectively, these observations are consistent with SNVs in driver lncRNAs being selected on the basis of functionally disrupting key elements, including secondary structures and RBP interaction sites.

### Mechanism for driver mutations via RBP-mediated stabilisation of oncogenic lncRNAs

RNA-binding proteins have the potential to stabilise or destabilise target RNAs. We hypothesised that driver SNVs in oncogenic lncRNAs may reduce the recruitment of destabilising RBPs, increasing lncRNA steady state levels and enhancing tumour fitness.

We focused on MALAT1, a well-characterised oncogenic lncRNA with numerous RBP interactions. SNVs in *MALAT1* target multiple binding sites for RBM22, DDX42, FTO and U2AF2 (Fig. 7a). For example, an FTO binding site at the *MALAT1* 3’ end, supported by eCLIP data and a high-confidence CNN-predicted motif containing a G->T SNV from a BRCA tumour that markedly reduced predicted binding affinity (Fig. 7b). Overall, six SNVs are predicted to alter FTO affinity for MALAT1, five of which decrease affinity (Fig. 7c).

**Figure 7:**
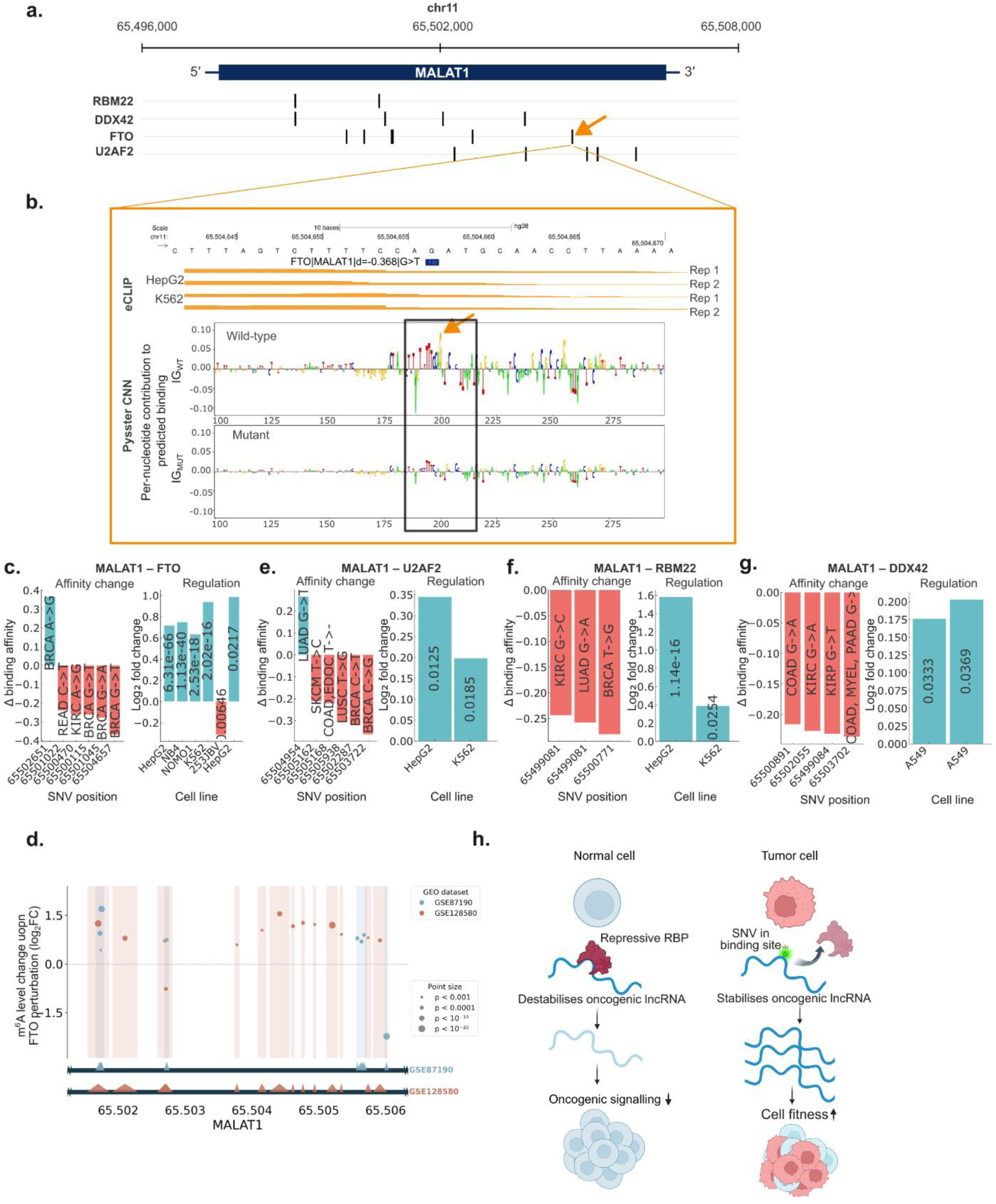
Model for driver SNVs via RPB-mediated stabilisation of oncogenic lncRNAs. a. Genomic organisation of MALAT1 onco-lncRNA showing eCLIP-defined binding sites for RBM22, DDX42, FTO and U2AF2. The FTO site indicated by the arrow is expanded in (b). b. An example of an SNV lying in the indicated FTO binding site in MALAT1. The location of the SNV is indicated by the filled rectangle. eCLIP read density maps are shown below for two replicates each in HepG2 and HeLa cells. Below, per-nucleotide integrated gradients (IG) attribution from the Pysster convolutional neural network (CNN) shows each base’s contribution to the predicted FTO-binding probability, for the wild-type (top) and mutant (bottom) sequence; letter height reflects the contribution, with positive and negative values indicating bases that increase or decrease predicted binding, respectively. c. Left panel: Impact of indicated SNVs in FTO eCLIP sites on Pysster-predicted binding affinity to MALAT1. The cancer type and nucleotide change are indicated inside bars. Right panel: Change in MALAT1 RNA levels as a result of FTO knockdown in indicated cell lines. Statistical significance (p-value) is indicated inside each bar. Expression data were obtained from GPSAdb database^64^. d. Change in m^6^A methylation in MALAT1 RNA resulting from FTO loss-of-function. *x*-axis denotes positions of detected m^6^A peaks along MALAT1 transcript, while the position of circles on *y*-axis indicates their log2 fold change in response to FTO loss-of-function. The colours and sizes of circles denotes their source (NCBI Gene Expression Omnibus identifiers) and statistical significance, respectively. The majority of points exhibit a positive change, indicating the FTO loss results in increased m^6^A methylation. (e–g). As for (c) but for indicated RBPs in MALAT1. h. Proposed model for driver mutations in lncRNAs: In healthy normal cells, recruitment of destabilising RBP suppresses onco-lncRNA activity. During tumourigenesis, driver SNVs disrupt RNA:RBP interactions, resulting in upregulation of onco-lncRNA activity and elevated cell pathogenicity and fitness.

To assess the impact of reduced FTO recruitment, we analysed RNA-seq data from a range of studies where FTO activity was knocked down by RNA interference^64^. FTO knockdown was associated with elevated MALAT1 levels in five out of six cell lines (Fig. 7c). This is consistent with the fact that FTO is an m^6^A demethylase and m^6^A methylation has been shown to promote MALAT1 oncogenic activity^65,66^. Further supporting this, knockdown of FTO leads to enhancement of m^6^A levels through the MALAT1 transcript (Fig. 7d). Thus, FTO is recruited to and destabilises the MALAT1 onco-lncRNA, and tumour SNVs tend to disrupt this interaction.

Similar patterns were observed for RBM22, DDX42 and U2AF2, each containing multiple affinity-reducing SNVs (including one recurrent) across different cancer types and consistent with upregulation of MALAT1 upon RBP loss (Fig. 7e,f,g).

This mechanism extends to other oncogenic lncRNAs. *SNHG3* and *SNHG14* harbour recurrent mutations in binding sites of destabilising RBPs (Fig. S7a-e). Conversely, multiple SNVs in NONO binding motifs of NEAT1 were predicted to increase the affinity between NONO and NEAT1 (5/6 cases), potentially enhancing paraspeckle formation and oncogenic activity (Fig. S7f,g).

Collectively, these data support a model in which driver SNVs target lncRNA:RBP interactions to promote tumourigenesis either by stabilising oncogenic lncRNAs (MALAT1, SNHG3, SNHG14) or enhancing their activity (NEAT1) (Fig. 7h).

## Discussion

The increasing scale of cancer genome sequencing is beginning to enable the detection of weak-effect and infrequently mutated non-coding drivers. Analysing whole-genome sequences from the 100kGP cohort, we define the largest catalogue of driver lncRNAs to date, comprising 121 genes under positive selection across 19 cancer types, approximately doubling existing catalogues. These include recurrently mutated oncogenes such as *NEAT1* and *MALAT1*, as well as many previously uncharacterised loci. Their disease relevance is consistently supported by diverse orthogonal evidence, including overlap with known cancer lncRNAs, germline susceptibility variants, mutational recurrence, transposon mutagenesis screens, and CRISPR perturbation data, collectively establishing lncRNAs as an important class of cancer driver genes. The relatively low overlap with previous studies indicates that many drivers remain undiscovered, consistent with the “weak driver” model in which numerous low-frequency mutations collectively shape tumour evolution^12^.

We report the first evidence that driver lncRNA mutations are mutually exclusive with established protein-coding drivers. This indicates that some driver lncRNAs are not independent of known oncogenic programmes but rather are integrated into those pathways. Such mutual exclusivity can also provide important mechanistic clues to elucidate the mechanisms of newly discovered driver lncRNAs.

These findings have translational implications. Many driver lncRNAs have independently been shown to modify disease when perturbed, making them candidates for emerging RNA-directed therapies^44^. Efforts are underway to therapeutically target oncogenic lncRNAs using antisense oligonucleotides and small molecules^44^. However, “cancer lncRNAs” are conventionally identified on the basis of correlative evidence such as tumour overexpression and validated in monolayer cell lines, leaving their true disease role and therapeutic potential uncertain. Positive selection on somatic driver mutations provides orthogonal and rigorous evidence for disease-promoting function, as demonstrated by the growing number of protein-coding drivers entering targeted therapy pipelines^67^. The present study identifies scores of lncRNAs with this level of support. Moreover, driver lncRNA mutations are present in a majority of tumours, including many lacking available targeted therapies, suggesting a large patient population could benefit from lncRNA-directed approaches.

A key advance of this work is the demonstration that sufficiently deep tumour genome data can be analysed akin to a high-resolution mutagenesis screen to map functional sites within lncRNAs. We show that driver mutations preferentially target conserved RNA secondary structures and RBP interaction sites. Computationally, these mutations are predicted to alter RNA folding energy and RBP affinity; experimentally, patient-derived variants induce structural rearrangements in vitro. Mutations cluster within core RBP recognition motifs and are enriched at the binding sites of cancer-associated RBPs, with a predominant effect of reducing binding affinity. Intriguingly, these effects may be mediated in part through the m^6^A epitranscriptomic pathway, linking RNA modifications to somatic driver mutations. From these observations, we derive a model in which somatic mutations modulate RBP recruitment - either disrupting repressive interactions or strengthening activating ones - to boost the activity of oncogenic lncRNAs. On the other hand, the impact of structure altering mutations is less clear, although there is recent precedent for disruption in lncRNA structures to promote oncogenesis^68^. These findings show how sufficiently large tumour genome cohorts can be used to simultaneously map disease genes and elucidate their molecular mechanisms, and opens the possibility of rationally designing therapeutic oligonucleotides that target specific lncRNA:RBP interactions.

Several limitations should be noted. Despite its scale, the 100kGP cohort does not saturate driver lncRNA discovery, as evidenced by the limited concordance across independent datasets and the absence of a plateau in downsampling analysis. Our element-level analyses depend on publicly available RBP binding maps, predominantly eCLIP data from K562 and HepG2, which may not capture the full diversity of RNA–protein interactions across cancer types. The 100kGP dataset also lacks matched RNA-sequencing data, requiring us to use TCGA for expression and survival analyses. These constraints are unlikely to alter our conclusions but will likely be addressed in larger future cohorts.

Taken together, this work defines a large, previously inaccessible class of cancer driver genes and provides the first mechanistic framework for how somatic mutations promote fitness by altering lncRNA function. These findings extend our understanding of the noncoding genome’s role in tumour initiation and evolution. The driver lncRNAs and their functional elements represent concrete therapeutic targets grounded in credible disease mechanisms. More broadly, we expect that the approaches developed here - for noncoding driver discovery and mechanistic characterisation - will be increasingly applicable as population-scale genome cohorts grow in number and diversity, forming the basis for a systematic understanding of noncoding RNA biology in cancer.

## Methods

### Ethics Statement and Consent

Whole-genome sequencing of matched tumour–normal samples was performed through the 100,000 Genomes Project (100kGP), approved by the East of England, Cambridge South Research Ethics Committee (REC reference: 14/EE/1112). All patients provided informed written consent, adhering to the Declaration of Helsinki (see https://www.genomicsengland.co.uk/initiatives/100000-genomes-project/documentation for further information on patient consent and withdrawal).

### Genome assembly

GRCh38 assembly was used for all analyses.

### LncRNA and protein-coding gene filtering

LncRNA annotations were obtained from GENCODE v39^34^. LncRNAs with any transcript having CPAT (Coding-Potential Assessment Tool) score >= 0.364 were excluded^69^. LncRNAs that overlap protein-coding genes on either strand were excluded. Furthermore, lncRNAs located within 10 kb of a protein-coding gene on the same strand were also removed. LncRNAs from the CLC3 set, sourced from Vancura et. al, were not subjected to these filters but were required to have a valid GENCODE identifier (648 / 702)^35^. Genes located within pseudoautosomal regions (PAR) and mitochondrial chromosome (chrM) were excluded from all analyses. These filters yielded a total of 7,794 lncRNAs for driver analysis (Supplementary File 1). Some lncRNAs analysed by ExInAtorPS were fully masked by blacklisted regions and were therefore excluded from downstream analysis (n = 7,163, Supplementary File 1).

### Blacklisted genomic regions

Problematic genomic regions were excluded by applying a comprehensive blacklist, constructed from the union of the following:

1. ENCODE basic blacklist (version 2, https://github.com/Boyle-Lab/Blacklist/tree/master/lists);
2. LINE-L1P regions (Genomics England) (https://re-docs.genomicsengland.co.uk/somAgg/);
3. Tandem simple repeats (Genomics England) (https://re-docs.genomicsengland.co.uk/somAgg/);
4. Segmental duplications (GIAB consortium): (https://ftp-trace.ncbi.nlm.nih.gov/ReferenceSamples/giab/release/genome-stratifications/);
5. Low mappability regions (GIAB consortium): (https://ftp-trace.ncbi.nlm.nih.gov/ReferenceSamples/giab/release/genome-stratifications/);

### Genome sample filtering

WGS data on 16,322 cancers were obtained from Genomics England data release version 17 (https://re-docs.genomicsengland.co.uk/release17/). These samples were cross referenced with the SomAgg v0.2 dataset. The following exclusion criteria were applied:

1. Samples with disease type “CHILDHOOD”, “OTHER”, “CARCINOMA_OF_UNKNOWN_PRIMARY”;
2. Samples from participants who did not provide data for use (as of 10.01.2024);
3. Samples whose karyotypic sex did not match the genomic sex;
4. Samples with more than 1% tumour cross-sample contamination;
5. Any samples whose match rank is not 1, 2, or 4; Genomics England defines various levels of “match rank” based on the corroboration between the genomic and the secondary clinical dataframes. Match rank of 3 and 5 imply little to no corroboration between genomic and clinical information (https://re-docs.genomicsengland.co.uk/cancer_analysis_histology/).
6. Samples were first separated by tumour type (primary, metastatic, and recurrent), retaining only the most recent sample per disease type for each participant. Samples from different disease types within the same participant were retained. After recombining the data, participants with multiple tumour types for the same disease were filtered down to a single representative sample using a strict hierarchy: Primary > Metastatic > Recurrent.
7. Remaining cancer types with <20 samples were excluded.
8. Tumours in colorectal (COAD, READ), stomach (STAD) and uterine cancers (UCEC, UCESC) were tested for microsatellite instability (MSI) using msisensor-pro (https://github.com/xjtu-omics/msisensor-pro) and those with a score of >3.5% were excluded^70,71^.

Following these filters, 12,631 samples from 37 cancer types remained.

### Filtering SNVs

Only variants that passed stringent quality control were retained. This included absence of sequencing or mapping artefacts as assessed using Genomics England internal filtering criteria (https://re-docs.genomicsengland.co.uk/somAgg/). Variants were required to be labelled “PASS” in the VCF files. The following variants were flagged and excluded from driver discovery analyses:

1. Germline allele frequency >1% in a Genomics England sub-cohort;
2. Germline allele frequency >1% in GnomAD;
3. Recurrent somatic variant with frequency > 5% in any Genomics England sub-cohort;
4. Overlapping tandem repeats;
5. Overlapping LINE1-LP regions;
6. Indels in homopolymer regions (>8 consecutive bases);
7. Common SNPs with allele frequency >= 1% in the 1000 Genomes project (annotated as ‘AF1000G’ in Genomics England VCF files (https://genome.ucsc.edu/cgi-bin/hgTrackUi?db=hg38&g=snp150Common).

### Identification of driver genes

ExInAtorPS was run with default parameters, considering only SNVs and indels of size 1bp. The previously described blacklisted regions were applied to all runs. The false discovery rate (FDR) was set at <= 10% for both protein-coding genes and lncRNAs. Functional impact of variants was assessed using CADD v1.6 scores (https://cadd.gs.washington.edu/)^72^.

As ground truth sets, the COSMIC Cancer Gene Census was used for protein coding genes and the Cancer LncRNA Census (CLC3) for lncRNAs^29,35^. A literature-curated list of potential false positive genes was obtained from Petrov et al^32^. One-sided Fisher’s exact test was used to calculate enrichment of true positives in predictions for both gene classes.

*XACT* (ENSG00000241743) was excluded from the candidate driver set due to a 300 kb+ long exon in GENCODE v39 annotation.

### Power analysis

Power calculations were carried out using the PCAWG power analysis scripts (https://github.com/broadinstitute/getzlabPCAWG-power_calculations)^17^. Background mutation rates for lncRNA were obtained from ExInAtorPS for each cancer cohort using mutation rates derived from flanking intergenic and intronic sequences. The median exonic length of lncRNAs (1155 bp), calculated from the GENCODE v39 annotation was used as the target size for power estimation.

### Simulation of mutations

Mutations were simulated using SigProfilerSimulator. Patient-specific tumour mutations were provided in the required format, and each cancer type was simulated independently. (https://github.com/AlexandrovLab/SigProfilerSimulator)^73^. A pan-cancer simulated set was generated by combining simulated data excluding lymphoid malignancies (LYMP) and melanoma (SKCM) samples. Default settings were used for all parameters (mutational signature contexts were set to ‘SBS96’ and ‘ID’). For empirical FDR estimation, 100 simulated sets were generated from breast cancer (BRCA) genomes. ExInAtorPS was applied to each set and the number of lncRNA drivers identified per simulated set was recorded. Empirical FDR was then estimated using:

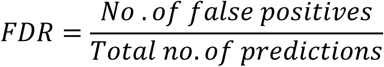

### LncRNA saturation analysis

Incremental subsets of the Breast invasive carcinoma (BRCA) cohort were generated, starting with 100 genomes and increasing in steps of 100 genomes per iteration. In total, 28 nested subsets were created, with the largest containing 2,800 patients. ExInAtorPS was applied to each subset, and the number of lncRNA drivers identified was recorded. Linear regression analysis was then performed in R using the lm function (number of drivers ∼ number of genomes), and the coefficient of determination (R²) was extracted from the model output.

### Mutational recurrence in lncRNAs

The recurrence ratio (RR) was defined as the number of mutations observed in the exon of a gene divided by the number of exonic mutated base positions. RR was calculated using the pan-cancer mutation dataset for both driver and non-driver lncRNAs. Samples from melanoma (SKCM), lymphoid malignancies (LYMP) and MSI tumours were excluded due to their localised hypermutation.

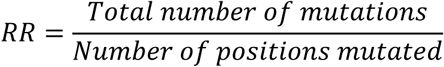

Analyses were restricted to non-redundant driver lncRNAs and non-driver lncRNAs. To assess whether mutations recur at specific positions beyond random expectation, we performed pan-cancer simulations preserving the observed trinucleotide context, thereby controlling for lncRNA length and tumour-type–specific mutational signatures. We quantified recurrence enrichment as the difference between the observed recurrence ratio and the mean recurrence ratio across 2000 simulated mutation sets. A one-sided Mann–Whitney U test was used to assess whether recurrence ratios were higher in drivers than in non-drivers.

### Germline single nucleotide polymorphism (SNP) enrichment

Cancer-associated SNPs were sourced from the EBI-GWAS catalogue (https://www.ebi.ac.uk/gwas/), and lncRNA annotations from GENCODE v39^34,74^. EBI GWAS catalogue SNPs were formatted for compatibility with bedtools, and multiple entries mapping to the same genomic locus were collapsed into unique entries. The germline SNP loci were then intersected with lncRNA loci and assigned to four categories: SNPs directly intersecting lncRNA exons, and those mapping within 1, 10, and 100 kb of the transcriptional start site (TSS). Germline SNP density (SNPs per Mb) was calculated, and log2 fold change of mean driver and non-driver SNP densities were computed across the four categories. Differences in SNP density distributions between the two groups were assessed using a one-sided Mann–Whitney U test for each category. Resulting p-values were adjusted for multiple testing using the Benjamini–Hochberg procedure.

### Enrichment analyses

CRISPR functional screens: Enrichment for “essential” or “hits” in cancer cell line CRISPR-inhibition screens compiled by Montero and Liu was assessed in driver lncRNAs^23,24^. Genomic coordinates were harmonised to the appropriate reference assembly (GRCh37 for Liu, GRCh38 for Montero, using GENCODE v47)^75^. LncRNAs were classified as hits if their TSS lay within 500 bp of a reported CRISPR screen hit. These were then annotated as driver or non-driver based on GENCODE IDs. Enrichment was evaluated using a one-sided Fisher’s Exact Test.

Transposon insertional mutagenesis (TIM) screens: Supplementary data containing lncRNA GENCODE gene IDs and corresponding TIM/Mb values was downloaded from the study by Esposito et. al^21^. Gene IDs were matched between the 100kGP driver set and TIM dataset. Enrichment was evaluated using a one-sided Fisher’s Exact Test.

Cancer lncRNA Census (CLC3): Supplementary data was obtained from Vancura et.al., comprising a list of GENCODE gene IDs and their associated features^35^. For disease-modifying CLC3 genes, we retained only those supported by evidence from in vitro or in vivo assays. For the analysis of oncogene and tumour suppressor proportions, we extracted annotations indicating whether each gene is classified as an oncogene, a tumour suppressor, or both, as reported in the supplementary dataset. Enrichment was evaluated using a one-sided Fisher’s Exact Test.

### Gene expression

Open-access TCGA (v41.0) RNA-seq gene expression data (transcripts per million, TPM units) and associated clinical survival data were downloaded from Xenahub (https://gdc.xenahubs.net)^76,77^. Matched normal tissue RNA-seq gene expression data (v10, TPM) were obtained from GTEx portal (https://gtexportal.org/home/downloads/adult-gtex/bulk_tissue_expression,GTEx_Analysis_v10_RNASeQCv2.4.2_gene_median_tpm.gct.gz)^78^.

Each TCGA cancer type was mapped to the most appropriate GTEx normal tissue for tumour–normal comparisons. 100kGP cancer codes were mapped to corresponding TCGA types; smaller 100KGP cohort subtypes without a clear TCGA match were excluded.

Differences in gene expression between groups was assessed using a one-sided Mann-Whitney U test.

### Survival

Overall survival (OS) was analysed using TCGA expression and clinical data with follow-up truncated at 5 years. For each gene, patients were stratified into high and low-expression groups based on the median. Cox proportional hazards models were fitted adjusting for age, sex and tumour stage.

Survival analyses by mutation status were performed using clinical data curated by Sosinsky et al^79^. Cancer-specific survival was performed following the authors’ published pipeline and dataset (Fig. 4; https://gitlab.com/genomicsengland/genomics_england_publications/100k_cancer_programme).

Analyses were conducted using the survival and survminer R packages (https://CRAN.R-project.org/package=survminer, https://CRAN.R-project.org/package=survival). Patients were stratified by presence or absence of mutations in each lncRNA. Univariate and multivariate Cox models were fitted, adjusting for age at diagnosis, sex, tumour stage and tumour mutational burden (TMB; log-transformed). Analyses were performed both within cancer types and pan-cancer. Pan-cancer models additionally included cancer type as a stratification factor to account for baseline hazard differences. Multiple testing correction was performed using the Benjamini–Hochberg method.

### Mutual exclusivity

Mutual exclusivity and co-occurrence were assessed using DISCOVER^80^. Binary alteration matrices (mutated/non-mutated) were constructed from WGS data for protein-coding genes and lncRNAs and analysed separately by cancer type. Protein-coding genes were classified as mutated if harbouring non-synonymous or loss-of-function variants (as defined by CellBase/ClinVar, https://re-docs.genomicsengland.co.uk/cancer_tiering/), whereas lncRNAs were considered mutated if containing ≥1 filtered exonic variant. Although DISCOVER inherently corrects for tumour mutation burden differences across samples and cancer-types, MSI and hypermutated samples were omitted from the analysis. Hypermutated samples were identified separately for each cancer type using the 90th percentile of the SNV count distribution as the threshold. Event matrices for protein-coding genes and lncRNAs were generated separately to account for differences in mutation rates and variant classes before testing for mutually exclusive or co-occurring pairs.

### ExInAtorEL

ExInAtorEL is a Python-based pipeline for identifying lncRNA elements enriched for somatic mutations using trinucleotide-aware subsampling. Inputs include: (i) tumour mutation catalogues (BED), (ii) lncRNA annotations (GTF) and (iii) element annotations (BED). Exonic regions are merged and filtered to define foreground (element-containing) and background (non-element exonic) regions. FASTA sequences are extracted and k-mer counts computed to enable trinucleotide-matched subsampling of background regions. Mutations are mapped to foreground (exonic elements) and background (exonic non-element) regions, and enrichment of each element assessed using a one-sided Fisher’s exact test. P-values are adjusted for multiple testing using the Benjamini–Hochberg procedure, with an FDR of < 0.1 considered significant. ExInAtorEL reports fold-enrichment values and adjusted p-values per element class, along with maps of mutated elements.

### RBP binding site identification using eCLIP datasets

High-confidence RBP sites were defined using ENCODE eCLIP data^55^. Peak files for each RBP in K562 and HepG2 cell lines were downloaded, and only those peaks replicated in both biological replicates retained. Overlapping peaks were merged to define consensus binding regions per RBP and cell line.

### RNA secondary structure analysis *in silico*

RNA secondary structures were predicted using RNAfold (ViennaRNA v2.0) under default parameters (37 °C). for paired wild-type and mutant sequences^59^. Minimum free energy (ΔG) was computed and mutation-induced effects quantified as ΔΔG = ΔG_mut − ΔG_WT. Absolute ΔΔG values were used to capture the magnitude of structural change.

### Mutations in RBP motifs

Motif instances within eCLIP peaks were identified using position weight matrices from CISBP-RNA (https://cisbp-rna.ccbr.utoronto.ca/), RBPDB (https://rbpdb.ccbr.utoronto.ca/), and DEWSeq using MOODS 1.9.4.1 (DNA mode; P < 0.005)^76,81–83^. Somatic mutations were mapped to the motif core and ±10 bp flanking regions.

Mutation enrichment in RBP motifs was assessed using ExInAtorEL. Motif regions were defined as foreground and the remainder of the exon as background.

The impact of mutations on RBP binding was predicted using the Pysster convolutional neural network^63^. For each mutation, a 400-bp window centered on the 5′ end of the overlapping eCLIP peak was extracted on the appropriate strand (reverse-complemented for minus-strand peaks). Pysster was used to predict the RBP-binding probability for the reference and alternative alleles, and the mutation impact score was defined as ΔP = P_mut − P_WT, the difference in predicted binding probability between the two alleles, where negative values indicate reduced binding and positive values increased binding. Per-nucleotide feature importance was visualised using integrated gradients. ΔP scores were compared between driver and non-driver lncRNAs using a one-sided Mann–Whitney U test with Benjamini–Hochberg correction (FDR < 10%).

### Changes in lncRNA expression upon RBP perturbation

Transcriptional responses of candidate lncRNAs to perturbation of their associated RBPs were obtained from GPSAdb v2.0 (https://gpsadb.com), a uniformly processed compendium of gene-perturbation RNA-seq datasets^64^. For each RBP of interest, perturbation datasets were identified using the Query module and filtered by cell line and perturbation modality (shRNA, siRNA, knockout). Pre-computed log₂ fold change values and adjusted p-values for the target lncRNA (perturbed versus control) were extracted directly from the GPSAdb differential expression tables and plotted without further processing.

### Differential m^6^A methylation upon FTO perturbation

Condition-specific changes in m^6^A methylation across candidate lncRNAs were retrieved from m6A-Atlas v2.0 (http://rnamd.org/m6a), which catalogues differentially methylated peaks called from MeRIP-seq experiments under defined perturbation conditions^84^. All available human FTO-perturbation datasets were queried; the direction of perturbation (knockdown or overexpression) was confirmed from the corresponding GEO records. For each significantly altered peak within a given lncRNA, the genomic coordinate (hg38) and log₂ fold change in m^6^A level were recorded. Peaks were plotted along the lncRNA locus, coloured by GEO accession, with point size scaled to statistical significance.

### RNA synthesis and purification

RNAs were synthesised using β-cyanoethyl phosphoramidite chemistry on a BioAutomation MerMade12 DNA/RNA synthesiser with RNA phosphoramidites (ChemGenes, GenePharma). RNAs were 5’-labelled with fluorescein (FAM) during synthesis. Deprotection was performed in aqueous ammonia/ethanol (3/1 v/v) for 16 h at 55°C. Silyl protecting groups were removed using triethylamine trihydrofluoride^85–87^. RNAs were desalted and purified by denaturing 12% PAGE.

### Native gel electrophoresis

FAM-labelled RNAs (5pmol) were folded in H_2_O by heating at 80°C for 3min followed by slow cooling to 50°C. At 50°C, 2x folding buffer was added, and samples incubated for 10min before slow cooling to 4°C. Final buffer conditions were either: (i) 100 mM NaCl, 3 mM MgCl_2_, 10 mM Tris-HCl pH 7.5 or (ii) 100 mM NaCl, 10 mM MgCl_2_, 10 mM Tris-HCl pH 7.5 Native loading buffer containing glycerol was added and samples were resolved on 10% native polyacrylamide gels at low voltage and visualised on a Amersham Typhoon Phosphorimager using MultiGauge Fujifilm software.

### Mapping of RNA structure with lead ion-induced cleavage

Lead ion-induced cleavage was performed on wild-type and mutated RNAs. Folded samples (5 or 4 pmol) in buffer (i) were mixed with lead acetate to final concentrations of 0.5, 1, and 2 mM Pb^2+^ and incubated for 15 min at 23°C. Control reactions received buffer only. Reactions were quenched with loading buffer containing 8 M urea and 10 mM EDTA and placed on ice. Cleavage products were resolved on 12% denaturing PAGE alongside formamide hydrolysis ladders (5mM MgCl_2_ 3.2 µl formamide for 15 or 30 min at 95°C) and T₁ RNase digestion ladders (50 mM sodium citrate pH 4.5, 8 M urea, 0.5U T₁ for 5 or 15 min at 55°C). Gels were imaged on an Amersham Typhoon Phosphorimager and analysed with MultiGauge Fujifilm software^62,88^. Background subtraction was performed and cleavage intensities normalised on a scale of 0 to 1.5. Strong cleavage was defined as ≥0.700, medium as 0.700–0.500. Results were compared to the prediction of the RNA model folding using by RNAstructure 4.6 software^60^.

## Acknowledgements

We wish to thank Andres Lanzos (Idunox), Roberta Esposito (University of Bern), Andrew Everall and Luis Zapata (Institute for Cancer Research), David Hughes, Thomas Lefeivre and Dylan Harvey (University College Dublin) for helpful advice and discussions. This work was supported by Research Ireland through Future Research Leaders award 18/FRL/6194 (R.J.), Consolidator Laureate award (IRCLA/2022/2500) (R.J.) and the Centre for Research Training in Genomic Data Science (S.R., R. Koya, J.R., B.D., S.B.). It was also supported by the National Science Centre of Poland through 2022/45/B/ST4/03586 (to R. Kierzek), 2021/41/B/NZ1/03819 and 2020/39/B/NZ1/03054 (to E.K.), 2020/39/D/NZ6/03267 (to M. S-J.), 2023/49/B/ST4/03763 (to M.S.). R.S.H. acknowledges grant support from Cancer Research UK (C1298/A8362) and the Wellcome Trust (214388). A.M. acknowledges SFB TRR267 (ID: 403584255, project Z02). We gratefully acknowledge the participants of the National Genomic Research Library (NGRL, National Genomic Research Library, Genomics England (2024) https://doi.org/10.6084/m9.figshare.4530893), whose contributions made this research possible. Secure access to the NGRL under project ID RR649 was provided by Genomics England, which delivers the NGRL in partnership with NHS England, and is wholly owned by the UK Department of Health and Social Care. The NGRL contains participants’ health data collected by the NHS as part of their care, along with samples and data from their participation in research, for which fully informed consent has been obtained. This includes genomic and clinical data provided through the NHS Genomic Medicine Service, as well as data obtained through research studies, including the 100,000 Genomes Project and the Generation Study, both of which are delivered in partnership with the NHS, and from other research cohorts involving external collaborators.

## Data Availability

Data from the National Genomic Research Library (NGRL) used in this research are available within the secure Genomics England Research Environment. Access to NGRL data is restricted to adhere to consent requirements and protect participant privacy. Data used in this research include:

For all the analyses performed in this paper we used NGRL dataset: somagg v0.2 and cancer analysis table V17.

All derived datasets and code for analysis can be found within /re_gecip/cancer_pan/ramnarayanan & re_gecip/cancer_pan/rkoya

Access to NGRL data is provided to approved researchers who are members of the Genomics England Research Network, subject to institutional access agreements and research project approval under participant-led governance. For more information on data access, visit: https://www.genomicsengland.co.uk/research

## Code Availability

ExInAtorPS: https://github.com/gold-lab/ExinatorPS [private repo]

## Supplementary Files

File 1: List of analysed lncRNAs

Description: GENCODE gene identifiers for all lncRNAs analysed by ExInAtorPS.

File 2: List of driver lncRNAs

Description: GENCODE gene identifiers for all driver lncRNAs along with associated cancer type.

File 3: Clinico-biological characteristics of driver lncRNAs

Clinical and biological features of driver lncRNAs with GENCODE gene identifiers and cancer type information.

File 4: Mutual exclusivity and co-occurrence analysis

Gene identifiers for significantly co-occurring and mutually exclusive lncRNA and protein-coding gene pairs.

File 5: Functional and regulatory elements within lncRNAs

Description: Summary of functional and regulatory elements within lncRNAs with data source.

File 6: ExInAtorEL enrichment analyses

Description: Cohort-wise mutation enrichment statistics for lncRNA elements, computed with ExinatorEL (one sheet per cohort).

File 7: List of RNA candidates shortlisted for experimental validation

Description: Summarised mutation impact, sequences, nucleotide variations, and associated CRS annotations for the shortlisted sites.

## Supplementary Figure Legends

**Figure S1:** a. The number of protein-coding driver genes predicted by ExInAtorPS in the 100kGP dataset. COSMIC Cancer Gene Census (CGC) genes are considered ground truth for driver genes (green). Known false positives taken from Petrov et. al.^32^ are coloured in orange. Remaining protein-coding genes are considered novel predictions. The proportions of these gene classes in the entire tested set of protein-coding genes is shown on the left for reference (‘EXPECTED’). Numbers inside bars represent gene counts, numbers above the bars indicate number of tumours in each cohort. b. The enrichment of true positive CGC driver genes amongst predictions in each 100kGP cancer cohort. Points represent indicated cancer types. All cancer types except Myeloma (MYEL) display significant enrichment at FDR≤10%. Statistical significance was determined by a one-sided Fisher’s exact test, with *p*-values adjusted for multiple testing using the Benjamini–Hochberg false discovery rate (FDR) method. c. The overall proportion of CGC genes amongst the union of predicted driver genes and tested non-driver genes (P=9e-109). Statistical significance was estimated using one-sided Fisher’s test. d. The proportion of known false positive (FP) driver genes from Petrov et. al.^32^ in the union of predicted driver genes and tested non-driver genes (P=0.8, one-sided Fisher’s test). e. The numbers of driver lncRNAs predicted (FDR≤10%) using real SNVs and shuffled SNVs for each cancer type. See Methods for details of SNV shuffling procedure. f. Histogram showing the numbers of driver lncRNAs predicted (FDR≤10%) in 100 different shuffled sets of BRCA SNVs.

**Figure S2:** a. Hazard ratios for MRPL20-DT in BRCA from a Cox proportional hazards model adjusted for age and tumour stage, corresponding to the Kaplan–Meier analysis shown in Fig. 4g. b. Hazard ratio for MALAT1 in BRCA adjusted for age, tumour stage and log₂(TMB), corresponding to the Kaplan–Meier analysis shown in Fig. 4h. c. Number of significant co-occurring gene pairs identified across all cancer types (no such gene pairs were identified outside of COAD and SARC cancer types). d. Mutually exclusive gene pairs in breast invasive carcinoma (BRCA) cohort at FDR≤1%.

**Figure S3:** a. SNV rates in elements within driver and non-driver lncRNAs. b. Coverage of driver lncRNA exons by indicated element classes. c. Number of significant driver elements identified across 100 sets of shuffled BRCA SNVs. No significant elements were detected in 96 of the 100 sets; in the remaining 4, the number identified was consistently low (n ≤ 27), far below the 158 elements identified from the real mutations. d. Driver element discovery in 14 driver lncRNAs identified from the PCAWG tumour dataset^21^. Here, pan-cancer SNVs from 100kGP were used. e. Identification of driver elements using shuffled SNVs from the indicated cancer types. Only driver lncRNAs corresponding to each cancer type were considered.

**Figure S4:** a. The evolutionarily conserved RNA structure (CRS) M0396124 within *EXOC3-AS1* lncRNA and tumour SNVs from 100kGP. Recurrently mutated positions are indicated by red boxes. b. The numbers of SNVs falling in CRS of driver lncRNAs that are predicted to reduce folding stability (Positive ΔΔG) or increase folding stability (Negative ΔΔG). Here, pan-cancer SNVs and the full set of driver lncRNAs are considered.

**Figure S5:** a, b. RNAfold-predicted secondary structures of wild-type (WT) and mutant (MUT) variants of conserved RNA structures (CRS) in driver lncRNAs *LINC01718* (a) and in *LINC01411* (b), showing substantial remodeling of the structure confirmed by lead ion (Pb²⁺) cleavage and increase in minimum free energy (ΔΔG). SNV locations are indicated by red circles. c, d. Native gel electrophoresis of in vitro synthesised RNA fragments of CRS from *LINC01718* (c) and in *LINC01411* (d). e, f. Lead ion (Pb²⁺) probing of the same in vitro synthesised fragments of *LINC01718* (e) and *LINC01411* (f). Note particularly the changes in Pb^2+^ accessibility of nucleotides in the red boxes, corresponding to predicted changes in base-pairing predicted by RNAfold above.

**Figure S6:** (a,b). Mutational enrichment of SNVs at RBP eCLIP-defined binding sites (x axis) versus sequence-motif sites within those binding regions (y axis), in the set of driver lncRNAs in BRCA and BLCA cohorts. Each point is one RBP. Only RBPs with significant enrichment in both tests are shown (eCLIP sites and motif sites, each FDR < 10%, tested independently using ExinatorEL). Dashed line, y = x (equal enrichment in both); points above indicate stronger enrichment at motif sites than across the full eCLIP region. (c-d). Mutation density across the binding motifs of RBP SRSF7 in pan-cancer (c) and BRCA (d) cohorts. The number of recorded SNVs is shown on y-axis at positions defined by the binding motif on the x-axis. The boundaries of the binding motif are denoted by vertical lines. The number of individual RBP site instances considered is indicated inside the box (‘Unique mutated-motif sites’). e. As for (a), but considering all RBPs and pan-cancer SNVs in non-driver lncRNAs. Note that no RBP type shows significant enrichment of SNVs in the core motif or eCLIP defined binding region at FDR < 0.1. f. Pysster-derived binding probabilities for the effect of a LIHC SNV in the RBP QKI site in *NEAT1*. g. Pysster-derived binding probabilities for the effect of a BRCA SNV in the RBP SRSF7 site in *MALAT1*.

**Figure S7:** a. Genomic overview of *SNHG3* showing eCLIP-defined binding sites for HNRNPK. b. Impact of SNVs on HNRNPK binding affinity to *SNHG3* (left) and up-regulation of the *SNHG3* level with HNRNPK knockdown in indicated cell lines (right). c. Genomic overview of *SNHG14* showing eCLIP-defined binding sites for SRSF7 and SRSF9. d. Impact of SNVs on SRSF7 binding affinity to *SNHG14* (left) and up-regulation of the *SNHG14* level with SRSF7 knockdown in HepG2 (right). e. Impact of SNVs on SRSF9 binding affinity to *SNHG14* (left) and up-regulation of the *SNHG14* level upon RPB knockdown in HepG2 (right). f. Genomic overview of *NEAT1* showing eCLIP-defined binding sites for NONO. g. Impact of SNVs on NONO binding affinity to *NEAT1*.

